# Game Theory Applications in Host-Microbe Interactions: *Mycobacterium tuberculosis* infection as an example

**DOI:** 10.1101/2019.12.26.888602

**Authors:** H Sharebiani, S Hajimiri, S Abbasnia, S Soleimanpour, H Asnaashari, N Valizadeh, M Derakhshan, R Pilpa, A Firouzeh, K Ghazvini, S Amel Jamehdar, SAR Rezaee

**Author notes:** **Corresponding author: SAR Rezaee** Immunology Research Center, Inflammation and Inflammatory Diseases Division, Medical School, Mashhad University of Medical Sciences, Mashhad, Iran. These authors contributed equally to this work (H Sharebiani, S Hajimiri).

## Abstract

The game theory describes the interactions between two players and the pay-off from wining, losing or compromising. *Mycobacterium tuberculosis* (*Mtb)*-host interactions were used as an example for the application of the game theory to describe and predict the possibilities of victory for any players. The gene expression for eight main markers of host response and three *Mtb* virulence factors were assessed in broncho-alveolar lavage of TB^+^ and TB^−^ patients. The game theory showed that a variety of paths exist that players can use, in response to the behaviour of the counterpart. Briefly, according to the “Nash equilibrium”, Ag85B is the main virulence factor for *Mtb* in active phase, however it is the most immunogenic factor if the host can respond by high expression of T-bet and iNOS. In this situation, *Mtb* can express high levels of ESAT-6 and CFP10 and change the game to the latency, in which host responses by medium expression of T-bet and iNOS and medium level of TGF-β and IDO. Consistently, The IDO expression was 134-times higher in TB^+^s than the TB^−^s, and the T-bet expression, ~200-times higher in the TB^−^s than the TB^+^s. Furthermore, *Mtb*-Ag85B had a strong positive association with CCR2, T-bet and iNOS, but had a negative correlation with IDO.

## Introduction

When two organisms, individuals, parties, teams, or even countries interact, the outcome varies greatly, depending on the intention of each party involved ^1^. Attempts to quantify these interactions can be assessed, using game theory models. Game theory is a mathematical framework, describing the outcome (pay-off) resulted from specific interactions (game) between two individuals (players) ^2,3^. In biologic context, game theory can describe interactions between a host and its parasite through epigenetic strategies, and the resulting pay-off from healthy, infected, or disease onset. The conflict is very complex, as both organisms are responsible for resisting and helping the associated genus to survive. In other words, host-microbe interactions are complex assemblies of large numbers of phenomena that are interacting competitively under multifactorial environmental conditions ^1–3^.

Bacteria and its host under stress may carry out the sophisticated principles of game theory, in order to decide whether to compromise or invade for elimination of the danger ^4^. Rationality of game theory indicates that each player in a particular game is motivated by maximizing his own pay-off. Therefore, it is especially important for players to think about each other’s strategic choice and react accordingly^4^. Interactions between *Mycobacterium tuberculosis* (*Mtb)* and human (players) are often included in the *Mtb’s* strategies to invade host responses, to replicate and persist within the host, and on the other hand, the host attempts to induce appropriate responses to eliminate the infection ^5^. Particularly, in case of *Mtb* infection these interactions are in a strict completion, as the microbe has adapted to replicate within the macrophages, which are specialized cells for killing microbes and the host must activate the infected cell to clear the *Mtb* ^5,6^.

The nature of *Mtb* and the genetic characteristics of the host seem to be important in the development, progress, and severity of infection. Even though, numerous association studies have been performed to identify genetic factors responsible for variations in TB susceptibility, to date, these candidate genes have not been found ^7,8^. Therefore, *Mtb* elimination and dormant or active *Mtb* infections in a given population seem to be associated with epigenetic processes. With a better understanding of the connections between bacterial infectious diseases and epigenetic events, opportunities will arise for therapeutic solutions, particularly, as epigenetic processes can be reverted. This opens a new pathway for future research in the field of microbial pathogenesis ^9,10^.

After infectious droplets of *Mtb* are inhaled, innate and adaptive effector mechanisms for intracellular infection or appropriate cell-mediated immunity (CMI) are able to eliminate the infection. The main effector player in this situation is the macrophages that are strongly activated by IFN-γ, and the habitats for *Mtb* transform into activated macrophages to eradicate the infection ^11^.

In an immunological equilibrium or latent TB, the main immune response against *Mtb* is CMI, which can only control the bacterial replication and consequent dissemination. A solid granuloma develops around the bacteria, composed of mononuclear immunity cells, such as macrophages in different maturation stages, and T cells of different phenotypes ^12^.

In reactivation or active TB infection, the weakening of the immune system or inappropriate immune responses causes the granuloma to become caseous and later to liquefy, and as a result, *Mtb* start to replicate. Then, they leave their host cells and spread to other areas of the lung, other organs, and the environment ^13^, that is called active TB.

In this study, the main elements in each phase of host immune responses and *Mtb* virulence factors were introduced in the game theory framework to show the application of this theory in host-microbe interactions. Briefly, the main strategies to adapt to its habitat, *Mtb*, as a parasitic organism, is to produce several virulence molecules, such as the 6-kD early secreted antigenic target ESAT-6 (EsxA), the 10-kD culture filtrate protein CFP-10 (EsxB), PPEs, and Ag85B, which actively intervene in both the innate and adaptive immune responses of the host ^14,15^.

Chemokines (CCLs) and chemokine receptors (CCRs) are the first factors, in response to a danger signal, involved in cell migration and immune polarization (such as Th1, Th2 and Th17). The polarization of these different types of responses depends on the APC functions, which can activate specific transcription factors like T-bet, GATA-3, ROR-γt, and FoxP3 toward polarization ^18,19^. For example, IDO activates in APCs catalyzes tryptophan and, thus, can induce a Treg response to produce immune-modulatory TGF-β, or APC- producing IL-12 can induce a Th1 response to secret IFN-γ ^20,21^. The last stage is activation of macrophages to produce oxygen-dependent, highly toxic factors for killing *Mtb*, proteases, inducing apoptosis or autophagy, or producing matrix metalloproteinases (MMPs), which may be involved in inflammation or tissue damages. These host activities may potentiate macrophages for killing the intracellular parasite or inhibit the response in favour of microbe disseminations ^22–24^.

In tuberculosis infection, *Mtb* and the host are two players in a game with a conflict of interest, in which interaction of their strategies determines the outcome ^25^. Among all possible strategies, appropriate responses of each player were selected by Nash equilibrium calculation as rational and optimal responses, happening in our assessments, and other possible strategies were excluded due to complexity. The interest of *Mtb* is to enhance its payoff by replicating as much as possible, and then by being transferred out of the host to find a new subject. In contrast, the host attempts to decrease or eliminate the infection with minimal harm ^25 25^.

In the present study, some main epigenetics factors were assessed and introduced to the game theory framework to describe the three outcomes of *Mtb-*host interactions, including host overcomes the infection (protection), equality (latency), or *Mtb* dissemination (active TB). Therefore, game theory was used as a mathematical model to describe the possibilities of those interactions.

## Methods

### Study population

The study population was 30 patients, including 14 TB patients and 16 non-tuberculosis pulmonary patients with positive Mantoux test results who referred to the Department of Internal Medicine, Ghaem Hospital, Mashhad University of Medical Sciences (MUMS), Mashhad, Iran. In the TB+ group, the smear and culture of sputum were negative, but IS6110-PCR on BAL sample was positive. For each patient, a clinical examination and a checklist were completed by a pulmonologist. The study was approved by the Bio-medical Ethics Committee of MUMS, (No: 941165, 930690, 930635 and 930634) and informed consent was obtained from each subject. In addition, all experiments were performed in accordance with relevant guidelines and regulations. In particular, written informed consent was obtained from all participants and all HIPAA identifiers were anonymized.

### Sample collection

A total of 50 ml pulmonary lavage was taken from each subject by a pulmonologist. After centrifugation at 7000 RPM, the pellet was stored in the Tri-pure (Roche Co., Germany) at −70 C°, until further RNA extraction.

### RNA extraction and cDNA synthesis

Total RNA was extracted from Tripure-treated samples (Roch Diagnostics Roche, Mannheim, Germany), according to the manufacturer’s instructions, and then reverse transcribed to cDNA with random hexamer primers, using the RevertAid™ H kit (Fermentas, Germany). All primers/probes for gene expression analysis were designed, using sequence data available in the NCBI databases (NCBI, NIH, USA) by Beacon Designer software (PREMIER Biosoft International, Palo Alto, CA). Specificity of the primers was checked by BLAST analysis (NCBI, NIH, USA). Primers and probes were synthesised by the BIONEER Corporation (South Korea), and their specificity was then confirmed by endpoint PCR, and sequencing of the products by Applied Biosystems (SEQLAB, Germany). Table 1 shows the sequences of primers and probes for the main virulence antigens of *Mtb* and the immunological factors in the main steps of the anti-responses *Mtb*.

**Table 1.**
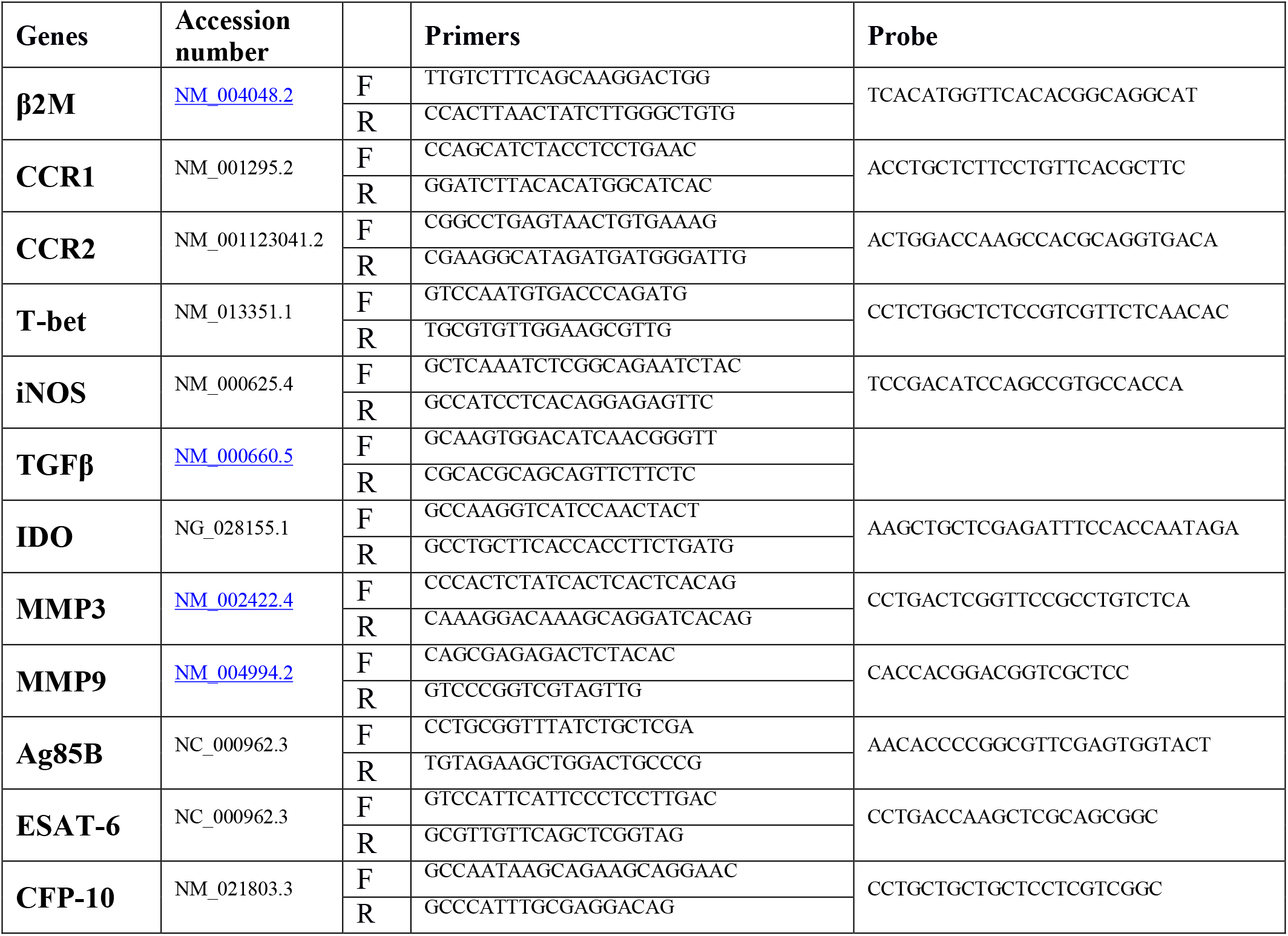
Primer and probe sequences & associated accession numbers.

### Relative real-time PCR

Quantitative TaqMan real-time PCR was performed with a Q6000 machine (Qiagen, Hilden, Germany), using TaqMan premix (Takara Corporation, Japan) as previously described ^39^. Six standards were prepared using 10-fold serial dilution of a concentrated sample of the genes of interest and a reference gene [beta-2 microglobulin (β2M)]. The normalized value of the expression for each gene was calculated as the ratio of the number of relative copies of the mRNA of interest to the number of relative mRNA copies of the reference gene, indicated as the expression index.

### Statistical Analysis

The Statistical Package for Social Sciences (SPSS) version 13 (IBM SPSS Inc., Baltimore, MD) was used for statistical analyses. Normality of the data was checked before data analysis using Kolmogorov-Smirnov test. For non-parametric analyses, the Mann-Whitney U test and Kruskal-Wallis test were used to compare gene expression between the two groups. The Spearman correlation coefficient was used to show statistical dependence or correlation between variables. Data was presented as mean ± SEM. Results were considered statistically significant if p ≤ 0.05.

### Game theory mathematical modeling of the host and *Mtb* interaction

The game theory was applied to the TB positive group, in which the main players in onset of tuberculosis are the host and *Mtb*. Therefore, their interactions can be modeled in two strategic (Tables 6 and 7) and extensive forms (Fig. 1) through backward induction. In all parts of the method, a dynamic game was assessed from the final (outcome) to the starting stages (initial interactions in infection). The players in this study used their intelligence in sequential attempts to reach a latency situation, which is in an equilibrium status with minimum risk and benefit; however, each player tried to win the game. The competition between the host and *Mtb* made the model a strictly competitive zero-sum game. To design this game, the first step was to determine two types of interactions: sequential and simultaneous. Therefore, a stage game was considered, in which the players simultaneously performed a static subgame with complete information during each stage while present within a dynamic game and sequentially responding to their opponents ^40–42^.

**Figure1:**
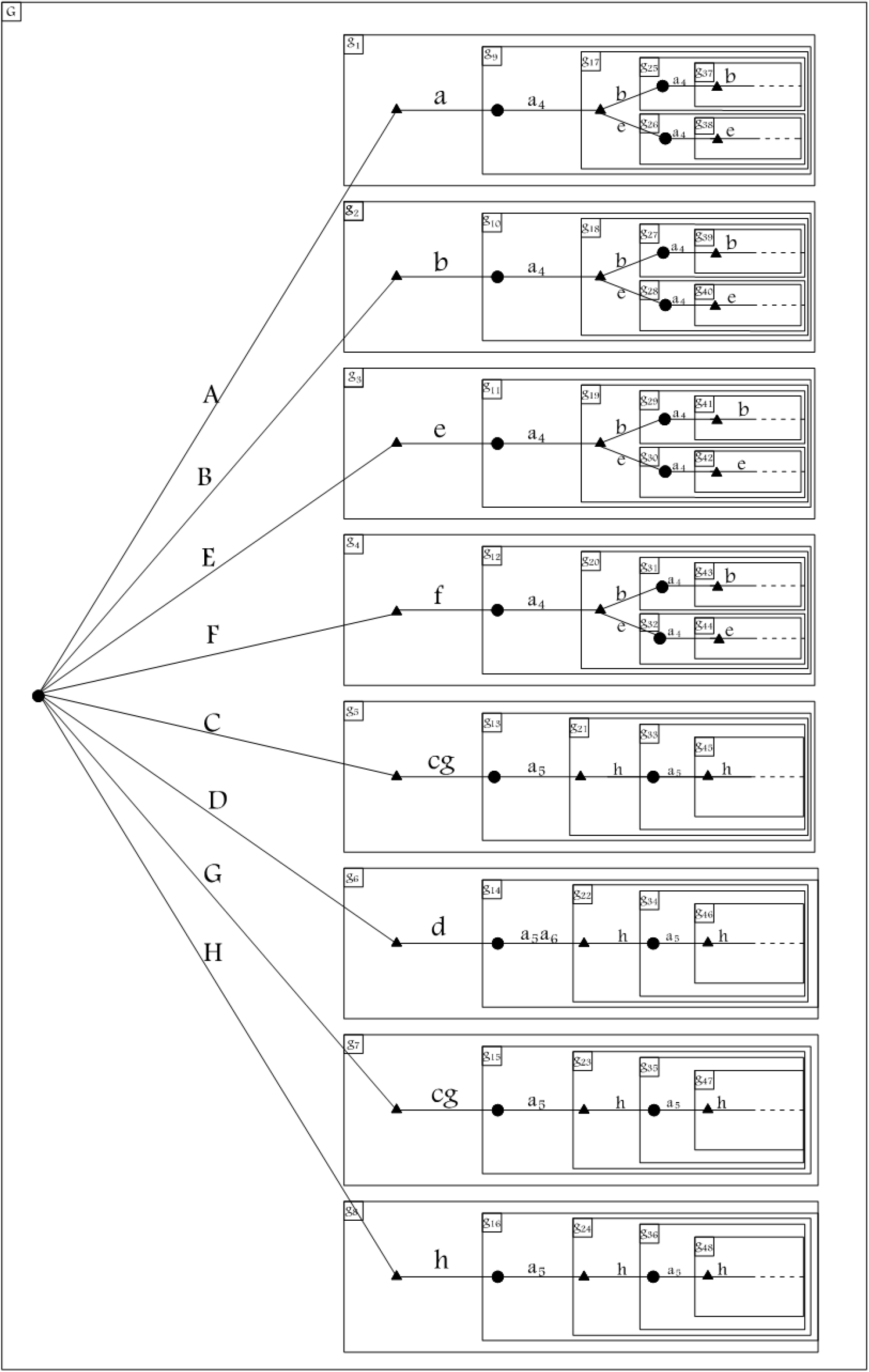
⚫ **Circle**; *Mtb’s* decision node.▲ **Triangle**; Host’s decision node.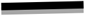 **Edges**; player’s strategy. …….. **Dotted lines**; Player’s frequently repeated strategies. **G**; Original game. **gn**; Subgame (which showed by rectangular nesting). **a_1_-a_6_**; *Mtb*’s actions. **A-H**; *Mtb*’s strategies (final outcomes). **a-h**; Host’s strategies. Subgame perfect equilibrium extensive form of game shows backward moving from repeated strategies (latency), which considered as the beginning of the game extended to the eight different *Mtb*’s strategies (A-H), by changing Ag85B, CFP-10 and ESAT-6 expression levels (reactivation or remaining latency). They are considered as final outcomes of host-*Mtb* interaction in each pathway. Host’s strategies “a”, “f” and “cg” are grim trigger strategies and “a_5_a_6_” is a grim trigger strategy performing by *Mtb* that host responded by strategy “d”. These strategies are reactivation strategies in favour of *Mtb* resulted in TB manifestation. Other *Mtb* and host strategies are cooperative strategies that are related to the latency phase.

### Quantification and design of the pay-off matrix

Since in a zero-sum game the pay-off for the *Mtb* is the negative of the pay-off for the host, each zero-sum game can be represented in a simplified form: a reward matrix. The matrix represents the pay-offs for the player who uses the strategies. The expression of main genes including CCR1, CCR2, T-bet, iNOS, TGF-β, Ag85B, CFP-10 and ESAT6 was considered as input data. Gene expression of selected genes have categorized (low/ high/ medium) by discretize by frequency operator and the pay-off matrix is designed based on conditional probability using the naive Bayes classifier (NBC) by Rapid Miner V5.3 software.

### Designing the extensive form of the game based on subgame Perfect Nash equilibrium and Pareto efficiency

The extensive form (or game tree) is a graphical representation of a sequential game. The game tree consists of nodes (or vertices), which are points at which players can take actions. They are connected by edges, which represent the actions that may be taken at the node. An initial (or root) node represents *Mtb*’s strategies (S_A_-S_H_) which begin the game due to entrance and invasion of bacteria and perform as a trigger for starting an interaction with host (game). Thus, pathways which define as every set of edges from the first node in the tree eventually arrives at the repetition stages, called A-H based on this origination (*Mtb*’s strategies (S_A_-S_H_)). Pathways A-H are consist of strategies that constitute a Nash equilibrium in every subgame of the original game, and this equilibrium is called the subgame perfect Nash equilibrium (SPE) ^40^. As explained earlier, the edges eventually arrive in repetition stages (shown as a dotted line in Figure 1) instead of a terminal node, because this is an infinitely repeated game, and the pay-off at the end of the game is calculated as a total pay-off and cannot be shown in the extensive form. In infinitely repeated games, the per-period interest rate (r) that results from game repetition is important, and players change the stage pay-off to the present value by a discount factor defined as 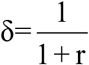. In order to calculate the total pay-off of an infinitely repeated game, a function was first defined as utility of strategy (*u*_*i*_ (*s*)) in each stage game (t) for every period (n) as follows:

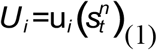

Then the players present value pay-off (V_i_) was calculated, and total pay-off was defined as the sum of the present values in each stage of the game for any time period calculated as:

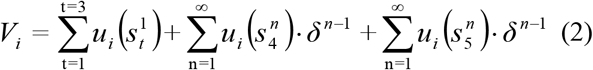

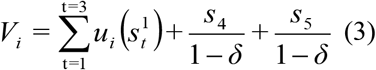

In infinitely repeated games, patience means is the increase of future pay-off in the present time; on the other hand, with an increase in repetitions, r goes to zero. Then, δ goes to one and future pay-offs resulting from a game repetition are preferred and more valuable than momentary pay-offs ^43^; this is called collusive pricing, which is a subgame Perfect Nash equilibrium outcome and leads to cooperation with Pareto dominant strategies ^41^.

### Parallel subgames’ Perfect Nash equilibrium and appropriate response efficiency

An equilibrium in which the player’s strategy constitutes Nash equilibrium in every parallel subgame set of the original game is a parallel subgame Perfect Nash equilibrium (PSPE), which is equal to the appropriate response, because it is determined by the maximization of the minimum pay-offs under consideration of all SPE strategies in a time period ^41^.

To determine the PSPE, the original game is separated into five parallel subgame sets, each of which refers to a stage. Nash equilibrium joint strategy pay-offs are placed on each set of a parallel subgame, and finally the best strategy of every parallel subgame is selected with the Min-Max method as PSPE. When a PSPE is found in any set of parallel subgames, the defined PSPE efficiency index is calculated for each path (A-H). The PSPE efficiency index is a ratio of the number of strategies that match the PSPE in past stages per the number of total strategies in past stages, which is represented as a percentage ^41^.

If “x” is the number of strategies which matched and “y” is the number of strategies which are unmatched with the PSPE in unrepeated parallel subgames. Also “n_0_” will be the number of unrepeated parallel subgames, so that n_0_=x+y ^41^.

If “n_r_” is the number of repeated parallel subgames that passed after the number of repetition times “t” and n_r_ = x′+y′, then “x′” is the number of strategies matched with the PSPE in repeated parallel subgames and “y’” is the number of strategies which are unmatched.

For paths repeated from the first stage, x′ /n_r_ represents the PSPE efficiency index in the repeated parallel subgames. In other paths, this index is represented by x/n_0_ calculated in unrepeated parallel subgames. Therefore, three types of strategies exist based on PSPE efficiency: PSPE dominant strategy (x>y or x′>y′), PSPE dominated strategy (x<y or x′<y′), and PSPE optimal strategy (x=y or x′=y′) ^41^.

The meaning of dominant and dominated strategies here differs from Pareto dominant and dominated, as this classification of strategies is based on PSPE efficiency, while classification in the previous parts was based on Pareto efficiency ^41^.

The PSPE efficiency index for each subject of paths repeated from the first stage (L(t) function) and other paths (R(t) function) are placed in a range of values based on the number of repetitions:

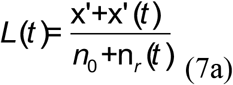

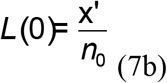

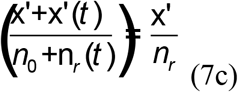

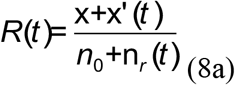

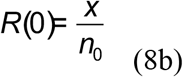

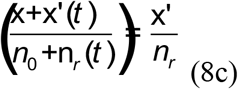

The PSPE efficiency index range for the path that earns “x” in unrepeated stages (n0) and “x′” in repeated stages (n_r_) is equal to [x/ n_0_, x′/n_r_), and the paths that are repeated from the first stage and earn “x′” have a constant value equal to x′/n_r_.

### Cooperative and grim trigger strategies in the original game

At the end of the modeling when all the last stages have been modeled, the game strategies are considered for the repeated stages and eight patient outcomes, and the earlier strategies that are placed in the repeated stages are cooperative strategies^44,45^. If one player contravenes his obligation, the first strategy that is not placed in the repeated stages is grim trigger strategies ^43^. Pay-off in a grim trigger strategy is less than that in cooperation strategies and could have different values of PSPE efficiency [x/ n_0_, x′/n_r_), while cooperative strategies are strategies with more pay-off that have a constant PSPE efficiency equal to x′/n_r_.

## Results

### Study population data

The mean age of TB^+^ patients was 68.43±8.29 years (range= 57 to 83 years), including 42.9% women. The mean age of TB^−^ controls was 69.56±7.46 years (range = 57 to 83 years), and 56.3% were males and 46% were females. However, the differences in age and gender distribution were not statistically significant (Table 2). Table 3 shows the social and demographic characteristics of the subjects studied. Table 4 shows the comparison of TB positive group with the TB negative group, in term of clinical and para-clinical symptoms.

**Table 2.**
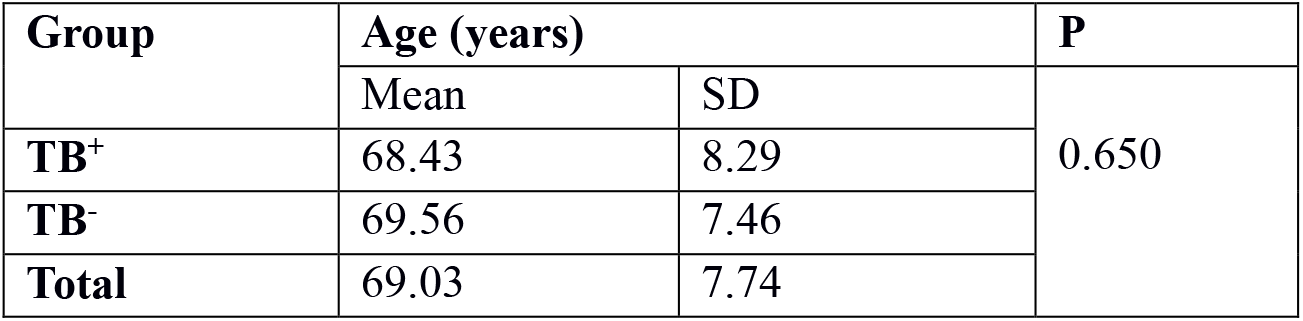
Age homogeneity of groups

**Table 3.** Social and demographic characteristics of the study subjects

| The socio-demographic profile |  |  | TB<sup>+</sup><br>N (%) | TB<sup>-</sup><br>N (%) | TOTAL<br>N (%) |
| --- | --- | --- | --- | --- | --- |
| Sex/ gender |  | Men | 57.1 | 43.7 | 50 |
|  |  | Women | 42.9 | 56.3 | 50 |
| Job | Men | Employee | 0 | 42.8 | 20 |
|  |  | Self-employed | 75 | 57.2 | 66.6 |
|  |  | Retired | 25 | 0 | 13.4 |
|  | Women | Housewife | 42.8 | 64.2 | 50 |
|  |  | Employee | 0 | 0 | 0 |

**Table 4.** Comparison of TB positive group with TB negative group in term of clinical and para-clinical symptoms

| Comparison of two groups in terms of clinical symptoms |  |  |  |  |  |  |  |  |  |
| --- | --- | --- | --- | --- | --- | --- | --- | --- | --- |
| Clinical Symptoms | TB<sup>-</sup> |  |  |  | TB<sup>+</sup> |  |  |  | P |
|  | YES |  | NO |  | YES |  | NO |  |  |
|  | Frequencies | % | Frequencies | % | Frequencies | % | Frequencies | % |  |
| Cough | 15 | 93.75 | 1 | 6.25 | 14 | 100 | 0 | 0 | 0.533 |
| Dyspnea | 14 | 87.50 | 16 | 12.50 | 11 | 78.57 | 3 | 21.43 | 0.433 |
| Lymphadenopathy | 0 | 0 | 16 | 100 | 0 | 0 | 14 | 100 | - |
| Weight loosing | 7 | 43.75 | 9 | 56.25 | 8 | 57.14 | 6 | 42.86 | 0.358 |
| Decreased appetite | 13 | 81.25 | 3 | 18.75 | 8 | 57.14 | 6 | 42.86 | 0.15 |
| Respiratory failure | 0 | 0 | 16 | 100 | 0 | 0 | 14 | 100 | - |
| Comparison of two groups in terms of para-clinical symptoms |  |  |  |  |  |  |  |  |  |
| Clinical symptoms | TB<sup>-</sup> |  |  |  | TB<sup>+</sup> |  |  |  | P |
|  | SD |  | Mean |  | SD |  | Mean |  |  |
| Albumin | 0.58 |  | 3.98 |  | 0.08 |  | 4.44 |  | 0.02 |
| LYM | 8.77 |  | 36.24 |  | 11.37 |  | 26.44 |  | 0.01 |
| PMN | 8.36 |  | 57.22 |  | 12.39 |  | 69.48 |  | 0.004 |
| WBC | 2.21 |  | 7.6 |  | 2.4 |  | 6.8 |  | 0.30 |
| ESR | 11.78 |  | 35.64 |  | 14.60 |  | 28.87 |  | 0.1 |
| CRP | 0.51 |  | 0.5 |  | 0.12 |  | 0.5 |  | 1 |

### The mRNA levels of CCR1, CCR2, iNOS, T-bet, TGFβ, IDO-1, MMP3, and MMP9 in the host

The CCR1 gene expression rates of BAL-PBMCs in TB^+^ and TB^−^ patients were 1.1±0.23 and 0.34± 0.09, respectively. There was a significant difference in the CCR1 gene expression between TB^+^ patients and TB^−^ controls *(p*<*0.01)*. Although, the average CCR2 gene expression in TB^+^ was around 3 times higher than the TB^−^ subjects, but no significant difference was observed in 95% confidence level. The CCR2 gene expression rates of BAL-PBMCs in TB^+^ and TB^−^ patients were 0.78±0.21 and 0.29±0.8, respectively *(p*<*0. 1*) (Table 5).

**Table 5.**
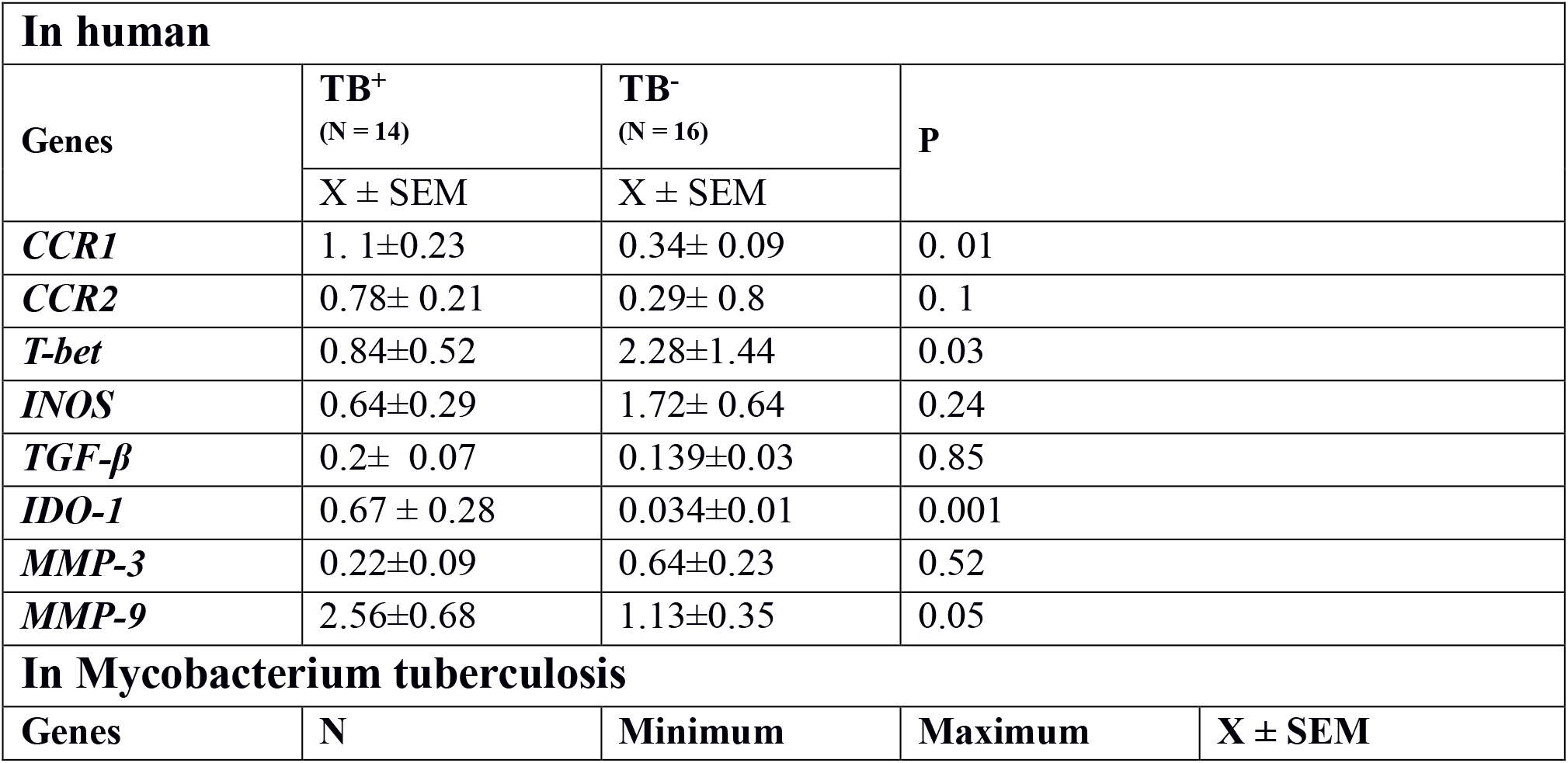
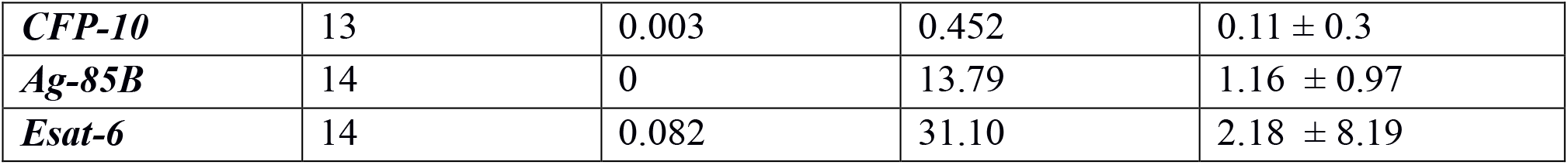
The gene expression data using Real-time PCR

**Table 6.**
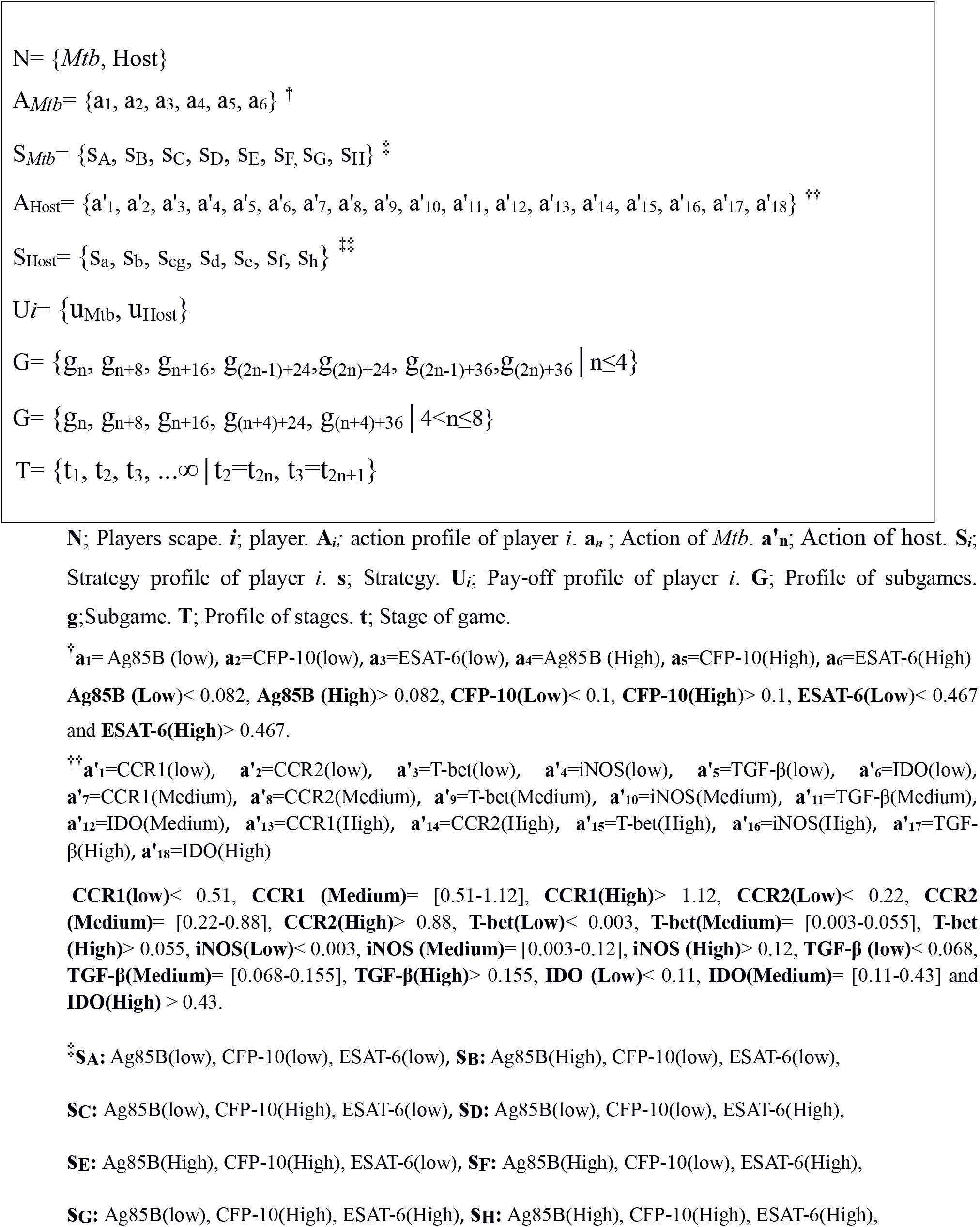
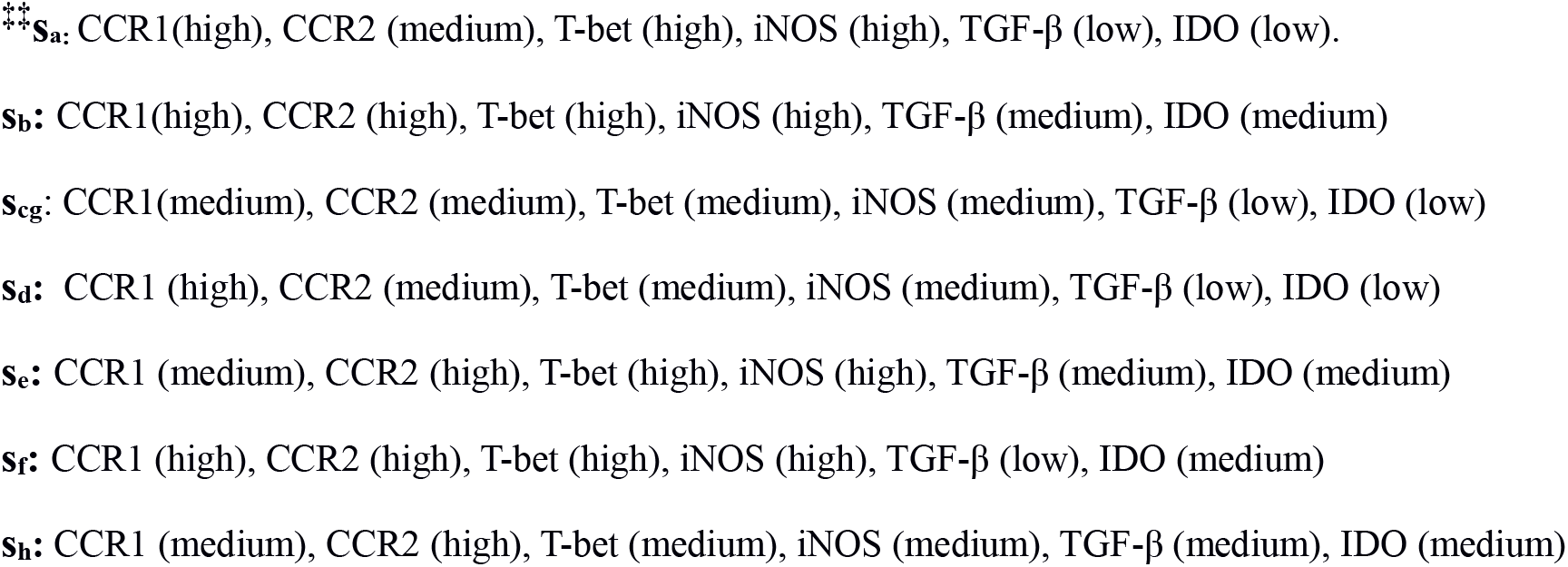
Strategic form of game.

**Table 7.**
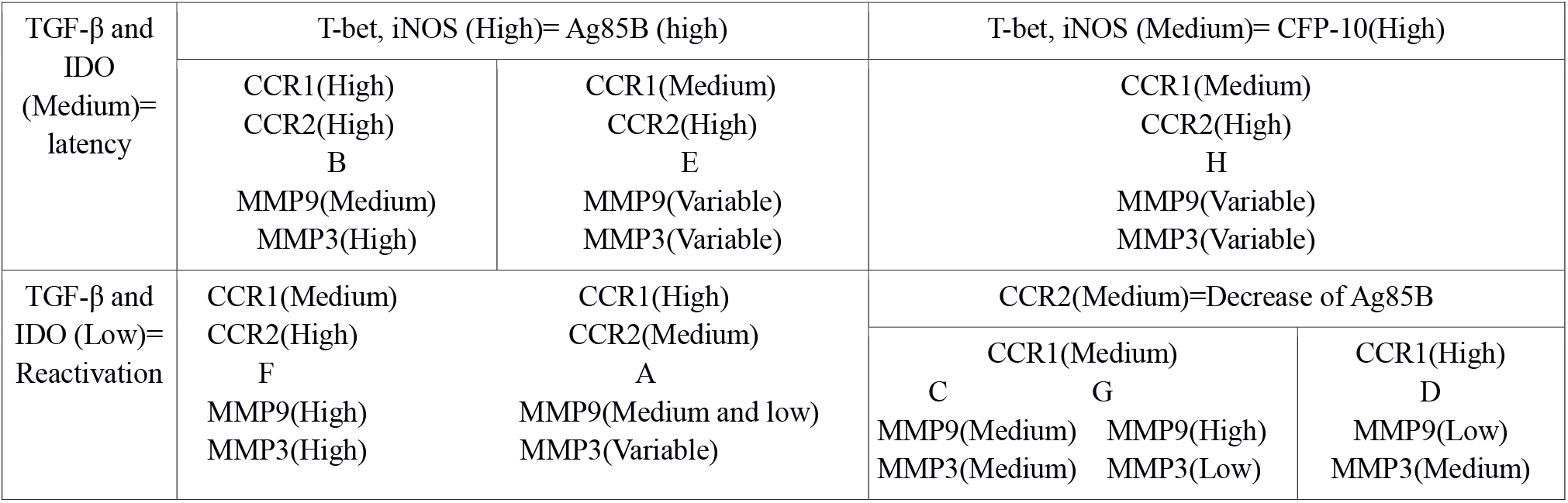
Evaluation of action roles in host strategies Expression of each gene in every host strategy was specialized in all strategies that remained latent (B, E and H paths), levels of TGF-β and IDO placed in medium range and in other paths that reactivated, these genes show low expression. Therefore, levels of TGF-β and IDO were differentiator between latent and reactivated paths. In all host strategies in response to high levels of Ag85B Mtb strategy, T-bet and iNOS expressed in high level while these levels were medium in response to high levels of CFP-10 or CFP-10 and ESAT-6, therefore levels of T-bet and iNOS expression related with type of Mtb strategies. Levels of CCR1 and CCR2 related to increase and decrease of Mtb virulence gene expression. High level of CCR1 expression was related to expression of CFP-10 and ESAT-6 in low level and medium level of this chemokine receptor had reverse effect on expression of these virulence genes. When CCR2 expressed in high level, expression of Ag85B placed on high level too, against in medium levels of CCR2 such as C, G and D paths, expression of Ag85B placed on low level.

The T-bet gene expression rates of BAL-PBMCs in TB^+^ and TB^−^ patients were 0.84±0.52 and 2.28±1.44, respectively; the difference was significant *(p*<*0.03)*. The average T-bet gene expression was around 1.53 times higher in TB^−^ patients than in the TB^+^ subjects.

The mean mRNA expression rates of iNOS in the BAL-PBMCs from TB^+^ and TB^−^ patients were 0.64±0.29 and 1.72± 0.64, respectively. As shown in Figure 1, mRNA expression was around 2.68 times higher in healthy carriers than in the TB^+^ patients; however, the difference was not statistically significant (Table 5).

The mean mRNA expression rates of TGFβ in the BAL-PBMCs of TB^+^ and TB^−^ patients were 0.2± 0.07 and 0.139±0.03, respectively. As shown in Figure 1, mRNA expression was around 0.69 times higher in TB^+^ patients than in the healthy carriers *(p*<*0.85).* However, no significant differences were found in the TGFβ mRNA levels between TB^+^ and TB^−^controls (Table 5) (Fig. 1).

The IDO-1 gene expression rates of BAL-PBMCs in TB^+^ and TB^−^ patients were 0.67±0.28 and 0.034±0.01, respectively. The average IDO-1 gene expression in TB^+^ was around 19.70 times higher than the TB^−^ subjects, and the difference was significant *(p*<*0.001)* (Table 5). The mean MMP3 mRNA expression rates in the BAL-PBMCs of TB^+^ and TB^−^ patients were 0.22±0.09 and 0.64±0.23, respectively, and the difference was not significant (Fig. 1) (Table 5). The MMP9 gene expression rates of BAL-PBMCs in TB^+^ and TB^−^ patients were 2.56±0.68 and 1.13±0.35, respectively. There was a significant difference in the MMP9 gene expression between TB^+^ patients and TB^−^ controls (*p*<*0.05*) (Table 5). The average MMP9 gene expression was around 2.26 times higher in TB^+^ patients than in the TB^−^ subjects.

### Extensive form of Game and Three Types of Cooperation

Figure 1 shows the extensive form of the game, in which the original game (G) is separated into subgames. Each path is an SPE of the original game. This is a Pareto dominant strategy, leading to cooperation and infinite repetition. There are three sets of strategies, each leading to specific repeated stages, consisting of a joint cooperative strategy. Every one of them shows one type of cooperation that equals latency strategies. The first set comprises the paths that begin with A, B, E, and F strategies and leads to the “e” host’s strategy, in response to a high level of Ag85B, which is *Mtb’s* strategy (a4, e). The second set comprises the paths that begin with A, B, E, and F strategies and leads to the “b” host’s strategy, in response to a high level of Ag85B, which is also *Mtb’s* strategy (a4, b). The third set comprises the paths that begin with C, D, G, and H strategies, and lead to the “h” host’s strategy, in response to a high level of CFP-10, which is *Mtb’s* strategy (a5, h). The D path, however, led to the “h” host’s strategy, in response to a high level of CFP-10 and ESAT-6, which is *Mtb’s* strategy (a5a6, h), before leading to the “h” host's strategy, in response to a high level of CFP-10, which is *Mtb’s* strategy (a5, h), just like the C, G, and H paths.

### Evaluation of Pathways by PSPE Efficiency as a Criterion for Appropriate Response

PSPE was determined between parallel subgames in each stage, consisted of “h” strategy in t5, a4 (Ag85B (High)) *Mtb* strategy in t4, the “h” host’s strategy against “a5a6” (high levels of CFP-10 and ESAT-6) *Mtb* strategy in stages t3 and t2, and finally in the paths that followed high levels of Ag85B (A, B, E, and F), the “a” host’s strategy, and in other paths that followed high levels of CFP-10 and ESAT-6 (C, G, D, and H), the “d” host’s strategy (Figure 1).

Figure 2 shows the PSPE efficiency indices calculated for each path. This index had a 50% constant value in the B, E, and H paths, classified into the PSPE optimal strategy (latent paths), 20% in the F, C, and G paths, and 40% in the A path, which is classified into the PSPE dominated strategy. The maximum value of this index was related to the D path (80%), which is classified into the PSPE dominant strategy.

**Figure 2:**
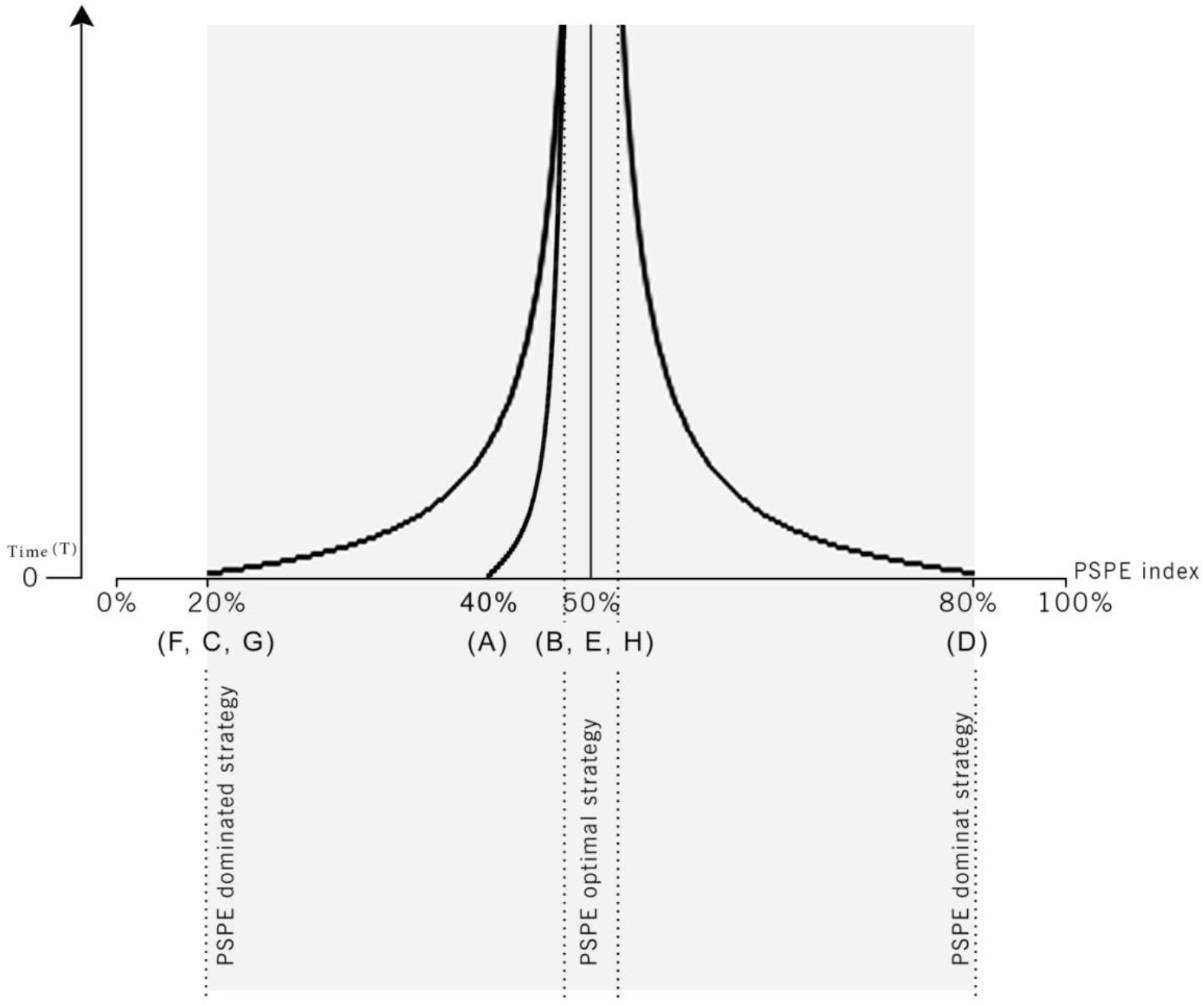
PSPE index during the time graph. Horizontal axis shows PSPE index from zero until 100% and vertical axis shows number of repetition times from zero until infinity. Right curve is a PSPE dominant strategy (D path), left curves are PSPE dominated strategy (A, F, C and G paths) and optimal strategy (B, E and H paths) is a middle line that have a constant value during the repetition times equal to 50% and this is vertical asymptote for two other curves. Increase of repeated time can lead to increasing of PSPE efficiency index and response appropriacy about inappropriate responses. These means that interaction between host and parasite will moderate by increase the number of encounters of players together and players avoid malicious strategies and move towards cooperation strategies like increase of immune response inhibition by medium level of TGF-β and IDO. Another argument is at last one of two players strategies which are in response to each other in repeated stages, must be placed in PSPE to take shape a cooperation strategy. In A, B, E and F paths Mtb strategy placed in PSPE and in C, D, G and H paths host strategy placed in PSPE.

Moreover, this value can be calculated in each repetition for each path, as figure 2 shows, in time course manner. Therefore, a long latency time could reduce the PSPE efficiency index to less than 80% in the D path, increase the PSPE efficiency index to higher than 20% in the F, C, and G paths, and increase it to more than 40% in the A path. Moreover, a long latency time is beneficial for the path that is classified into the PSPE dominated strategy, and a short latency time is beneficial for the path that is classified into the PSPE dominant strategy (Fig. 2).

### Evaluation of the pathways by the level of MMPs expression

To evaluate pulmonary tissue damage as a patient’s outcome of *Mtb*, the host interaction levels of MMP9 and MMP3 were determined and categorized into levels of low, medium, or high. Levels of MMP3 and MMP9 were determined in each path from A to H. The results showed that in the D path, the MMP9 level was classified into the low range; in the A path, it was classified into the low and medium ranges; the B and C paths were related to medium levels of MMP9, and the G and F paths showed high levels of MMP9. Paths E and H had variable values. Paths B and F were classified into high levels of MMP3; the D and C paths were classified into the medium range; the G path showed low levels of MMP3, and the H path was placed in the medium and low ranges. Paths E and A had variable values.

### Reactivation or Grim Trigger Strategy

In the A, C, F, and G paths, the host contravened his obligation in the t1 stage and entered a grim trigger strategy, classified as a PSPE dominated strategy. The pay-off decreased and the PSPE efficiency index diminished from 50% to 20% for the C, F, and G paths and from 50% to 40% for the A path. Therefore, reactivation was performed by inappropriate responses in C, F, and G, and finally led to a bad outcome (high levels of MMP9 in G and F and medium levels in C). In the A path, reactivation performed at a 40% efficiency. This is twice that of the C, G, and F paths, even though it was placed on the PSPE dominated strategy. Yet, as a result, this path showed a better outcome with medium and low levels of MMP9.

In the D path, *Mtb* contravened his obligation (high levels of CFP-10 expression as a cooperative strategy) in the t2 stage, and entered a grim trigger strategy (high levels of ESAT-6 expression in addition to CFP-10), classifying as a PSPE dominant strategy. Because of this switching, pay-off decreased, and the PSPE efficiency index increased from 50% to 80%. Therefore, reactivation was performed by appropriate responses that led to better outcomes (low levels of MMP9).

According to the results, the A, F, C, and G paths represented suitable strategies in latency that were reactive by the host and led to bad outcomes, while the D path represented a suitable strategy in latency that was reactive by *Mtb* and finally led to a good prognosis. In the B, E, and H paths, no player contravened from his obligation in any of the t1 or t2 stages. Therefore, the players remained in their joint cooperative strategies with a constant pay-off and PSPE efficiency index (latent group).

## Discussion

In this study, the status of host-microbe interactions (game) in the onset of TB (win or lose) was investigated in active and latent stages of infection.

Therefore, to use game theory, we had to choose suitable variables, which are implicated in microbe and host strategy by evaluation of gene expression (epigenetics outcome) for each player (*Mtb* and host) in each subject (TB^+^), simultaneously. Therefore, the gene expression of the main virulence factor of *Mtb* (Ag85B,CFP-10 and ESAT-6) were considered as *Mtb*’s strategy, and the host’s main immunological molecules were assessed in four different steps of immune responses as host’s strategies: (i) leukocyte recruitment by chemokines receptors (CCR1 and CCR2), (ii) polarization of the response by transcription factor expressions (T-bet and TGF-β), (iii) cytokine and immunomodulatory factors production (IDO and iNOS), and release of effector molecules in tissue damage (MMP3 and MMP9). However, this condition can be changed depending on host (human) or microbe (*Mtb*) activities, under the influence of epigenetic strategies ^26^. In the present study, the first step of the game was induced by *Mtb* as a parasite for colonisation in which the host respond by the recruitment of leukocytes to the site of infection. Findings showed that both CCR2 and CCR1 expression were higher in TB^+^ patients than in Mantoux positive TB^−^ patients. Although, the CCR1 expression was statistically significant, the CCR2 was only in level of 90% confidence interval met significance (0.1). This form of expression is in favour of bacteria as Th2-like cells and monocyte macrophages recruitment and exacerbate airway inflammation.

The next step was consistent with this findings as the main difference between TB^+^ and TB^−^ patients was the expression of T-bet, which was ~200 times more in the Mantoux positive TB^−^ patients than in the active TB patients. Furthermore, the expression of IDO as an immune-suppressor was 134 times more in TB^+^ than in the TB^−^ patients. In such situation, with a high level of IDO expression as an inducing factor for Treg polarization, a higher expression of TGF-β in TB^+^ patients is expected, while in reality, its expression was 4 times more, but did not meet the 95% CI (P=0.1). Taken together, Th1 recruitment (T-bet high expression) can protect the host against *Mtb* infection and Th2 like, Treg high responses and may be the CCR2 expressing monocyte/macrophage have been in favor of lung inflammation and microbe dissemination in TB^+^ patients.

The expression profile for the TB^+^ patients were applied in a mathematical model to describe players’ behaviours in the game, and by the Pareto efficiency criteria to describe the three outcomes of *Mtb* infection, in terms of host-microbe interactions: the colonisation of *Mtb* to establish and disseminate infection, (Pareto dominated and excluded), clearance by the host immune response and host overcomes the infection (protection, Pareto dominated and excluded), equality (latency, Pareto dominant and selected). As a result, the discussion formed around the Nash equilibrium in a type of Pareto dominant strategies that led to latency. In this model, an extensive form of the game was designed by SPE to exclude non-credible threats that a rational player would not actually carry out, because it would not be in his interests ^27^. Therefore, none of the players selected low levels of defective factors or virulence molecules as a strategy, although these strategies were also not seen among the players, in the assessed situations.

In this study, it could be assumed that all patients (TB^+^ and TB^−^) were in the latent phase, based on the mean age of the subjects and positive Monteux test results. At this stage of latency, the interactions of host and microbes reach a cooperation strategy or latency status, but some subjects experienced *Mtb* reactivation or grim trigger strategy. Some patients followed the strategies, in response to high levels of *Mtb* Ag85B expression (b and e strategies; Fig. 1), and others designed different strategies in response to high levels of *Mtb* CFP-10 expression (h strategy; Fig. 1).

The best strategy for *Mtb* in latency is the high level of Ag85B expression that was determined to be a component of PSPE. This can be explained by the fact that Ag85B is a TAG synthase and might be a key player in both cell wall stability and the biosynthesis of storage compounds for the survival of *Mtb,* in the dormant state (*Mtb* in the A, B, E, and F paths followed this strategy) (Fig. 1).

Another strategy that *Mtb* may use in latency is high levels of CFP-10 expression. In this stage, all ESAT-6 molecules were in the form of CFP-10/ESAT-6 as a chaperone and were not able to express any effective function (The C, D, G and H paths followed this strategy). As a result, high levels of CFP-10 expression and the heterodimer form of CFP-10/ESAT-6 guarantee the compromise between the host and *Mtb*, due to (i) inhibition of phagosome rupture and cytosolic translocation of mycobacteria that prevent growth, proliferation, and the dispersion of bacteria; (ii) ESAT-6 in complex with CFP-10 also interacts with beta-2 microglobulin (β2M), affecting the antigen presentation activities of macrophages and prevents the recognition of *Mtb* in dormancy; (iii) CFP-10 in the ESAT-6/CFP-10 complex may particularly contribute to neutrophil recruitment and activation. Thus, during *Mtb* infection, the high level of this virulence factor leads to inflammatory reactions, complicating the immuno-pathogenesis of TB ^28^; (iv) ESAT-6/CFP-10 complex enhances the production of NO and IL-12 released from M1 cells, following IFN-γ stimulation. The game theory results for this status showed that the presence of CFP-10 resulted in the production of medium level T-bet, inducing Th1 cells to produce IFN-γ, and also medium level of iNOS for NO production ^7,28^. These two mechanisms keep the pathogen in a limited activity (latency); and (v) In such strategy, the high level of CCR2 expression may induce IFN-γR1 on the surfaces of the macrophages and create an appropriate innate immune response to help with *Mtb* elimination ^18^.

In pathway C and G, when the host contravenes his obligation from “h” to “cg” strategy, lead to reactivation by a grim trigger strategy (cg), instead of a cooperative strategy (h). Furthermore, the production of IDO and TGF-β was decreased, in order to up-regulate the protective immune response to remove *Mtb*, also the probability of phagosome acidification increased, and CFP-10 and ESAT-6 were dissociated upon acidification. Consequently, ESAT-6 allowed to interact with the phagosomal membrane, resulting in phagosome rupture and cytosolic translocation of mycobacteria. Another altered part of the host strategy is the decrement of CCR2 expression, leading to the decreased recruitment of monocytes. This is a rational decision, because ESAT-6 was shown to directly bind to Toll-like receptor 2 (TLR2), inhibiting TLR signaling in macrophages. Inhibition of macrophage response by ESAT-6 does not earn a significant pay-off ^29^. Therefore, the recruitment of monocytes will be decreased in the host, in order to prevent the pay-off of energy loss. This is a rational way to minimize *Mtb* pay-off by not allowing the bacteria to develop its own strategy.

In pathway D, the patient induces a pattern of gene expressions, including CCR1 (high), CCR2, T-bet, iNOS, TGF-β, and IDO (medium), in response to *Mtb’s* high expression of CFP-10 in latent phase. In this commensal situation, *Mtb* decides to leave the cooperation and expresses ESAT-6 along CFP-10, leading to the increased probability of ESAT-6 production without CFP-10. Therefore, these ESAT-6 molecules can inhibit phagosome maturation and lead to phagosome rupture and cytosolic translocation of mycobacteria. With this strategy, bacteria can evade the toxic elements of the phagolysosome, and it can activate pro-inflammatory pathways such as IL-1, IL-8, IL-12, and IFN-β. Thus, this is an appropriate host immune response to low levels of TGF-β and IDO, high levels of CCR1, and medium levels of CCR2. That is a punishment strategy for *Mtb,* because of (i) low levels of TGF-β and IDO lead to a decrease in immune system over-reaction by inhibiting harmful inflammation; (ii) change of CCR1 from the medium to the high level, consequently led to decreased CFP-10, as activation of this pathway recruits Th1 lymphocytes to the site of infection, which are the most powerful arm of the immune system to combat intracellular invaders. The radical change in CCR2 from a high to a medium level did not allow Th2 to come to the site of the infection ^30^. Consequently, the response promotes in favour of Th1, and the host can produce appropriate anti-*Mtb* responses. Thus, *Mtb* loses the game, and has to decrease its production of the main virulence factor, Ag85B, due to the high level of immunogenicity. Finally, *Mtb* could just save a high level of ESAT-6, because the virulence is weak for dominance; it is a punishment state for *Mtb*. Moreover, low levels of MMP9 and medium levels of MMP3 in this path, show a low level of pulmonary tissue damage, in favour of a modulated proper immune response.

Low levels of MMP9 lead to increased recruitment of neutrophils and an appropriate innate immune response, while medium levels of MMP3 can be related to the medium level of CCL2 (MCP-1), as a ligand of CCR2 and to medium levels of monocytes recruitment ^31–34^.

The game theory model demonstrated that the best outcome in path “D”, occurred for several reasons. The first one is related to the Pareto efficiency, like the other paths, but with only one difference, which is 80% of its strategies were matched with PSPE and appropriate protective responses. The rationale behind the reason that this strategy is better for the host is, unlike the other paths, in path D *Mtb* leaves the cooperation or latency by a grim trigger strategy, which is matched with PSPE. Firstly, *Mtb* is a parasite and the most common behavior for any parasite is to “save its host”, in favour of species survival. Reaching an equilibrium state with the host (latency) is the best strategy for *Mtb*, and perhaps for the host.

If *Mtb* changes the strategy to a very severe infection, not only the host loses the game and dies due to the severity of infection, but also the parasite loses its ecological niche, which is not in favour of the survival of the species. Therefore, adaptation of the host and the parasite by cooperation strategy for survival is rational as game theory showed by PSPE optimal paths B, E and H at 50% efficiency. On the other hand, the parasite must have an evasion strategy not to lose the game. Thus, as the results of the current study showed, *Mtb* never changed strategy from high levels of CFP-10 to high levels of Ag85B. Although, Ag85B is a powerful virulence factor for the bacteria, it is also an immune-dominant antigen for the host, and induces a severe response to eliminate the bacteria. As a result, high levels of ESAT-6 and CFP-10 were the best decision to replace with high levels of CFP-10, because ESAT-6 is a weaker virulence molecule than both the Ag85B and CFP-10, and even an immune-modulator for host Th1 responses ^35,36^.

The second response is the host’s response to high levels of ESAT-6 and CFP-10, differing from latency strategies, in term of TGF-β and IDO levels. Furthermore, it differs from other reactivation strategies, in response to high levels of CFP-10, high levels of CCR1, and medium levels of CCR2 ^36^.

Among the paths (A, C, G, and F), in which the host contravenes his obligation and the new strategy is PSPE dominated, the path A reactivation strategy was more efficient and comparable with path D. In fact, A and D paths are analog, which are appropriate responses to high levels of Ag85B and CFP10-ESAT-6, respectively.

In pathway A, the *Mtb* virulence factors in the lower levels and host strategies are a, b and e, which can be frequently repeated in b and e. As can be concluded from these figures, *Mtb* first starts the colonization, and the host induces a higher expression of CCR1 and CCR2, T-bet, and iNOS, and IDO, and TGF-β expression rates are medium. In such situation, *Mtb* tries to induce high expression of Ag85B (a4) frequently, until it reaches to a host strategy (“a”), in which CCR1, T-bet, and iNOS remain at high levels, but CCR2 is at a medium level, and TGF-β and IDO are at a lower level. In these levels, the patients leave the cooperative strategy or latency, and *Mtb* can win the game with a higher expression of Ag85B, in which the players enter, a grim trigger strategy with 40% PSPE efficiency, which is 10% less than the cooperative strategy. Therefore, it should be noted that the host leaving a commensal interaction, results in a better situation for *Mtb* to escape from the host’s immune responses and enter reactivation status. It should be noted that the host’s strategies in these paths, only differ in the levels of T-bet and iNOS; however, the TGF-β and IDO levels remain unchanged, which is in favour of *Mtb* reactivation.

The lower levels of IDO and TGF-β can be in favour of the host, in clearing the infection. As previously mentioned, IL-10 and TGF-β are also important anti-inflammatory cytokines, modulating immune responses ^16^. Appropriate amounts of TGF-β and IL-10 (like medium level in latency strategies) in the *Mtb*-infected organs are vital, for balancing the Th1 protective response and preventing Th1 hypersensitivity (DTH) reactions, by suppressing the intensity of this response to bacteria. Thus, in *Mtb* infection, the importance of IL-10 and TGF-β may be double-edged ^16^. High expression of TGF-β and IDO is a non-credible strategy that is excluded in this model, since either dampening the host’s inflammatory response or limiting the Th1 appropriate response goes toward immuno-compromise, which may allow reactivation and dissemination of infection to an active TB or reactivation of a latent status. According to the current results, it can be expected that, in the presence of high levels of IDO expression, as an inducing factor for Treg polarization, TGF-β production will also increase significantly, but in gene expression data, this assumption did not meet this status (CI=90%, p=0.1). Of note, patients in this study had no immuno-compromising conditions ^16^.

Another change in the host response is related to the CCRs expression that leads to changes in cell recruitment of Th1, Th2, Treg, and Th17 for clearance, infection establishment, or granuloma formation ^37^. Besides physically sequestering mycobacteria, the formation of granuloma inhibits bacterial growth by subjecting mycobacteria to stressful condition, such as starvation, reactive oxygen, nitrogen intermediates, and hypoxia. However, viable, apparently extracellular bacteria can persist in chronic granulomas for many years ^37^. In a suitable status like immune-compromised host, *Mtb* can overcome the pressure of the host’s immune system and be reactivated. As this model shows, there are direct associations between the strength of virulence factor, Ag85B, CFP-10 and ESAT-6, and the host’s immune response, such as CCR-1, T-bet, Th1 polarization, IFN-γ production and iNOS, for macrophage activation.

The potentiation of the innate immune response in this model was determined by IFN-γ and iNOS, leading to NO production, and was responsible for the potentiation of the phagocytes for elimination of pathogens. As mentioned, the iNOS expression level is directly related to the severity of virulence factors, regardless of latency or reactivation phase, like T-bet expression. It seems that this is a “tit-for-tat’’ strategy, in which one player responds in a stage with the same action that his opponent used in the last stage.

For the last stage of the effector phase, MMPs assessment is the outcome of the responses in clearance or pulmonary tissue damage, which can be discussed in the context of the pathways C, G, and D. It should be noted that MMPs are also able to modify chemokine and cytokine activity, resulting in modified inflammatory cell recruitment ^24,38^.

The immune response in both paths C and G had the same high levels of CFP-10, but the expression of MMPs differs. In path C, the medium level of MMPs contributes to the remaining *Mtb* infection. In path G, in the presence of the low level of MMP3, the CCL2 degradation decreased, providing more CCL2 ligands for the medium expression of CCR2 (same levels as in paths C and G than the path C). Therefore, monocytes are recruited more to the inflamed site of infection in the G path than to the path C and do not allow Th2 to come. Consequently, the response is in favour of Th1, and the host can produce a strong appropriate reaction. On the other hand, a high level of MMP9 causes damage to pulmonary tissue, and by increasing degradation of IL-8, it decreases neutrophils recruitment at the infection site. In this strategy, the host does not have an appropriate innate immune response. Finally, in path G, high levels of ESAT-6-CFP-10 induce this outcome. Low levels of MMP9 and the medium expression of MMP3 contribute to an appropriate innate response, in producing a moderate Th1 response and monocyte recruitment.

The study have some limitations, the life and its maintenance is one of the most complicated phenomenon in the universe, and science could not find a reliable methodology for study of the complexity. Particularly, “the conflicts on survival in parasitism” in which behavior of two living players in the site of interactions must be studied, simultaneously. It is nearly impossible to include all of the factors which are implicated in the site of microbe-host and micro-environment interactions in a three dimensional system. Therefore, to introduce the game theory as a model for prediction of the outcome of such interactions, only main *Mtb* and host activities were included.

## Conclusion

According to the “Nash equilibrium” although Ag85B is the main virulence factor of *Mtb* for proliferation and winning the game, the most immunogenic factor arises when the host can response by high expression of T-bet and iNOS and defeats the microbe. In such situation of immune response, *Mtb* can express high expression of ESAT-6 and CFP10 and drives the game to a host strategy with a medium expression of T-bet and iNOS. In addition to these host factors, it is more likely that TGF-β and IDO are differentiating factors between latent and reactive phases, which their medium expression levels can lead to the latency and low levels to the reactivation.

Accordingly, it can be assumed that effective vaccines, containing Ag85B and maybe ESAT6 (without suppressive C-terminal) should be considered in different attempts to protect the host from *Mtb* infection. Furthermore, with the respect of host responses, the application of the appropriate adjuvants and the assessment of expression of T-bet and maybe CCR2 in PBMCs, in the presence of the multi-stage vaccine cocktail will be in favour of Th1-response. Taken together, the present study as an example of application of the game theory along with epigenetics assessments, open a new path to study the microbe/ tumor cell and host immune response, simultaneously with the onset of the disease and therapy.

## Acknowledgements

Many thanks to the Vice Chancellor of Research, MUMS, Mashhad, Iran for supporting the studies for compailing this article, Biomedical Eethics committees number of 941165, 930605, 930634 and 930690. These studies are the subjects of student thesis.

## Author Contributions

RS and SH designed the study. SH, HS, AS, VN, PR, FA and AH performed the experiments and analyzed the data. RS, SH and SS wrote the paper. AH as a pulmonoliest took the lavage, All authors have read and approved the fnal version of the manuscript.

## Additional Information

## References

1 von Neumann J, M. O. Theory of games and economic behaviour. Princeton, NJ: Princeton University Press. (1944).

2 Schmidt, E. F. a. K. M. A Theory of Fairness, Competition, and Cooperation. he Quarterly Journal of Economics 114, 817–868 (1999).

3 Lambert, G., Vyawahare, S. & Austin, R. H. Bacteria and game theory: the rise and fall of cooperation in spatially heterogeneous environments. Interface focus 4, 20140029, doi:10.1098/rsfs.2014.0029 (2014).

4 Tago, D. & Meyer, D. F. Economic Game Theory to Model the Attenuation of Virulence of an Obligate Intracellular Bacterium. Frontiers in Cellular and Infection Microbiology 6, 86, doi:10.3389/fcimb.2016.00086 (2016).

5 Ehrt, S., Rhee, K. & Schnappinger, D. Mycobacterial Genes Essential for the Pathogen’s Survival in the Host. Immunological reviews 264, 319–326, doi:10.1111/imr.12256 (2015).

6 Gengenbacher, M. & Kaufmann, S. H. E. Mycobacterium tuberculosis: Success through dormancy. Fems Microbiology Reviews 36, 514–532, doi:10.1111/j.1574-6976.2012.00331.x (2012).

7 Peddireddy, V., Doddam, S. N. & Ahmed, N. Mycobacterial Dormancy Systems and Host Responses in Tuberculosis. Frontiers in Immunology 8, 84, doi:10.3389/fimmu.2017.00084 (2017).

8 Cheepsattayakorn, A. & Cheepsattayakorn, R. Human genetic influence on susceptibility of tuberculosis: from infection to disease. Journal of the Medical Association of Thailand = Chotmaihet thangphaet 92, 136–141 (2009).

9 Kathirvel, M. & Mahadevan, S. The role of epigenetics in tuberculosis infection. Epigenomics 8, 537–549, doi:10.2217/epi.16.1 (2016).

10 Esterhuyse, M. M., Linhart, H. G. & Kaufmann, S. H. Can the battle against tuberculosis gain from epigenetic research? Trends in microbiology 20, 220–226, doi:10.1016/j.tim.2012.03.002 (2012).

11 Khademi, F., Derakhshan, M., Yousefi-Avarvand, A., Tafaghodi, M. & Soleimanpour, S. Multi-stage subunit vaccines against Mycobacterium tuberculosis: an alternative to the BCG vaccine or a BCG-prime boost? Expert review of vaccines 17, 31–44, doi:10.1080/14760584.2018.1406309 (2018).

12 Ahmad, S. Pathogenesis, Immunology, and Diagnosis of Latent Mycobacterium tuberculosis Infection. Clinical and Developmental Immunology 2011, 814943, doi:10.1155/2011/814943 (2011).

13 Lenaerts, A., Barry, C. E. & Dartois, V. Heterogeneity in tuberculosis pathology, microenvironments and therapeutic responses. Immunological Reviews 264, 288–307, doi:10.1111/imr.12252 (2015).

14 Tang, X. L. et al. CFP10 and ESAT6 aptamers as effective Mycobacterial antigen diagnostic reagents. The Journal of infection 69, 569–580, doi:10.1016/j.jinf.2014.05.015 (2014).

15 Forrellad, M. A. et al. Virulence factors of the Mycobacterium tuberculosis complex. Virulence 4, 3–66, doi:10.4161/viru.22329 (2013).

16 Soleimanpour, S. et al. APC targeting enhances immunogenicity of a novel multistage Fc-fusion tuberculosis vaccine in mice. Applied microbiology and biotechnology 99, 10467–10480, doi:10.1007/s00253-015-6952-z (2015).

17 Mosavat, A. et al. Fused Mycobacterium tuberculosis multi-stage immunogens with an Fc-delivery system as a promising approach for the development of a tuberculosis vaccine. Infection, genetics and evolution : journal of molecular epidemiology and evolutionary genetics in infectious diseases 39, 163–172, doi:10.1016/j.meegid.2016.01.027 (2016).

18 Domingo-Gonzalez, R., Prince, O., Cooper, A. & Khader, S. Cytokines and Chemokines in Mycobacterium tuberculosis infection. Microbiology spectrum 4, 10.1128/microbiolspec.TBTB1122-0018-2016, doi:10.1128/microbiolspec.TBTB2-0018-2016 (2016).

19 da Silva, M. V. et al. Expression Pattern of Transcription Factors and Intracellular Cytokines Reveals That Clinically Cured Tuberculosis Is Accompanied by an Increase in Mycobacterium-Specific Th1, Th2, and Th17 Cells. BioMed Research International 2015, 591237, doi:10.1155/2015/591237 (2015).

20 Blumenthal, A. et al. M. tuberculosis induces potent activation of IDO-1, but this is not essential for the immunological control of infection. PloS one 7, e37314, doi:10.1371/journal.pone.0037314 (2012).

21 Boer, M. C., Joosten, S. A. & Ottenhoff, T. H. M. Regulatory T-Cells at the Interface between Human Host and Pathogens in Infectious Diseases and Vaccination. Frontiers in Immunology 6, 217, doi:10.3389/fimmu.2015.00217 (2015).

22 Rivera-Marrero, C. A. et al. M. tuberculosis induction of matrix metalloproteinase-9: the role of mannose and receptor-mediated mechanisms. American journal of physiology. Lung cellular and molecular physiology 282, L546–555, doi:10.1152/ajplung.00175.2001 (2002).

23 Lam, A. et al. Role of apoptosis and autophagy in tuberculosis. American journal of physiology. Lung cellular and molecular physiology 313, L218–l229, doi:10.1152/ajplung.00162.2017 (2017).

24 Ong, C. W., Elkington, P. T. & Friedland, J. S. Tuberculosis, pulmonary cavitation, and matrix metalloproteinases. American journal of respiratory and critical care medicine 190, 9–18, doi:10.1164/rccm.201311-2106PP (2014).

25 Eswarappa, S. M. Location of pathogenic bacteria during persistent infections: insights from an analysis using game theory. PloS one 4, e5383, doi:10.1371/journal.pone.0005383 (2009).

26 Ip, M. et al. Human epigenetic alterations in Mycobacterium tuberculosis infection: a novel platform to eavesdrop interactions between M tuberculosis and host immunity. Hong Kong medical journal = Xianggang yi xue za zhi 21 Suppl 7, S31–35 (2015).

27 Heifetz, A. Game Theory Interactive Strategies in Economics and Management. (Cambridge University Press, 2012).

28 Esmail, H., Barry, C. E., Young, D. B. & Wilkinson, R. J. The ongoing challenge of latent tuberculosis. Philosophical Transactions of the Royal Society B: Biological Sciences 369, 20130437, doi:10.1098/rstb.2013.0437 (2014).

29 Pathak, S. K. et al. Direct extracellular interaction between the early secreted antigen ESAT-6 of Mycobacterium tuberculosis and TLR2 inhibits TLR signaling in macrophages. Nature immunology 8, 610–618, doi:10.1038/ni1468 (2007).

30 Hirsch, C. S., Rojas, R., Wu, M. & Toossi, Z. Mycobacterium tuberculosis Induces Expansion of Foxp3 Positive CD4 T-cells with a Regulatory Profile in Tuberculin Non-sensitized Healthy Subjects: Implications for Effective Immunization against TB. Journal of clinical & cellular immunology 7, 428, doi:10.4172/2155-9899.1000428 (2016).

31 Rodriguez, D., Morrison, C. J. & Overall, C. M. Matrix metalloproteinases: what do they not do? New substrates and biological roles identified by murine models and proteomics. Biochimica et biophysica acta 1803, 39–54, doi:10.1016/j.bbamcr.2009.09.015 (2010).

32 Salgame, P. MMPs in tuberculosis: granuloma creators and tissue destroyers. The Journal of Clinical Investigation 121, 1686–1688, doi:10.1172/JCI57423 (2011).

33 Moores, R. C., Brilha, S., Schutgens, F., Elkington, P. T. & Friedland, J. S. Epigenetic Regulation of Matrix Metalloproteinase-1 and −3 Expression in Mycobacterium tuberculosis Infection. Front Immunol 8, 602, doi:10.3389/fimmu.2017.00602 (2017).

34 Shi, C. & Pamer, E. G. Monocyte recruitment during infection and inflammation. Nature reviews. Immunology 11, 762–774, doi:10.1038/nri3070 (2011).

35 Karbalaei Zadeh Babaki, M., Soleimanpour, S. & Rezaee, S. A. Antigen 85 complex as a powerful Mycobacterium tuberculosis immunogene: Biology, immune-pathogenicity, applications in diagnosis, and vaccine design. Microbial pathogenesis 112, 20–29, doi:10.1016/j.micpath.2017.08.040 (2017).

36 Guo, S. et al. The CFP10/ESAT6 complex of Mycobacterium tuberculosis may function as a regulator of macrophage cell death at different stages of tuberculosis infection. Medical hypotheses 78, 389–392, doi:10.1016/j.mehy.2011.11.022 (2012).

37 Flynn, J. L., Chan, J. & Lin, P. L. Macrophages and control of granulomatous inflammation in tuberculosis. Mucosal Immunology 4, 271–278, doi:10.1038/mi.2011.14 (2011).

38 Brew, K., Dinakarpandian, D. & Nagase, H. Tissue inhibitors of metalloproteinases: evolution, structure and function. Biochimica et biophysica acta 1477, 267–283 (2000).

39 Kianmehr, M. et al. Immunomodulatory effect of characterized extract of Zataria multiflora on Th1, Th2 and Th17 in normal and Th2 polarization state. Food and chemical toxicology : an international journal published for the British Industrial Biological Research Association 99, 119–127, doi:10.1016/j.fct.2016.11.019 (2017).

40 Indrajit Raya, S. S. Observable implications of Nash and subgame-perfect behavior in extensive games. Journal of Mathematical Economics 49, 471–477 (2013).

41 Nash, J. Non-Cooperative Games. Annals of Mathematics 54, 286–295 (1951).

42 Sorin, S. Chapter 4 Repeated games with complete information. Handbook of Game Theory with Economic Applications 1, 71–107, doi:https://doi.org/10.1016/S1574-0005(05)80007-4 (1992).

43 Barton L. Lipman, R. W. Switching costs in infinitely repeated games. Games and Economic Behavior 66, 292–314, doi:https://doi.org/10.1016/j.geb.2008.04.018 (2009).

44 Smith, J. M. The theory of games and the evolution of animal conflicts. Journal of theoretical biology 47, 209–221 (1974).

45 R Axelrod, W. H. The evolution of cooperation. Science 211, 1390–1396, doi:10.1126/science.7466396 (1981).

