## appendix for "Game Theory Applications in Host-Microbe Interactions: *Mycobacterium tuberculosis* infection as an example"

**Backward induction:** It is an iterative process for solving finite extensive form or sequential games. First, one determines the optimal strategy of the player who makes the last move of the game. Then, the optimal action of the next-to-last moving player is determined taking the last player's action as given. The process continues in this way backwards in time until all players' actions have been determined. Effectively, one determines the Nash equilibrium of each subgame of the original game.

**Cooperative game:** It is the game in which players are able to make enforceable contracts. Hence, it is not defined as games in which players actually do cooperate, but as games in which any cooperation is enforceable by an outside party (e.g., a judge, police, etc.). In termed non-cooperative games, contracts must be self-enforcing.

**Dominant strategy:** A strategy is dominant if, regardless of what any other players do, the strategy earns a player a larger payoff than any other. Hence, a strategy is dominant if it is always better than any other strategy, for any profile of other players' actions. If one strategy is dominant, than all others are dominated.

**Dominated strategy:** A strategy is dominated if, regardless of what any other players do, the strategy earns a player a smaller payoff than some other strategy. Hence, a strategy is dominated if it is always better to play some other strategy, regardless of what opponents may do. If a player has a dominant strategy than all others are dominated, but the converse is not always true.

**Dynamic game:** When players interact by playing a similar stage game numerous times, the game is called a dynamic, or repeated game. Unlike simultaneous games, players have at least some information about the strategies chosen on others and thus may contingent their play on past moves.

**Equilibrium:** An equilibrium, (or Nash equilibrium, named after John Nash) is a set of strategies, one for each player, such that no player has incentive to unilaterally change her action. Players are in equilibrium if a change in strategies by any one of them would lead that player to earn less than if she remained with her current strategy.

**Extensive form:** The extensive form (also called a game tree) is a graphical representation of a sequential game. It provides information about the players, payoffs, strategies, and the order of moves. The game tree consists of nodes (or vertices), which are points at which players can take actions, connected by edges, which represent the actions that may be taken at that node. An initial (or root) node represents the first decision to be made. Every set of edges from the first node through the tree eventually arrives at a terminal node, representing an end to the game. Each terminal node is labeled with the payoffs earned by each player if the game ends at that node.

**Game:** The interaction among rational players and the decisions of some players impacts the payoffs of others. A game is described by its players, each player's strategies, and the resulting payoffs from each outcome. In sequential games, the game stipulates the timing (or order) of moves.

**Grim trigger strategy**: A trigger strategy usually applied to repeated prisoner's dilemmas in which a player begins by cooperating in the first period, and continues to cooperate until a single defection by her opponent, following which, the player defects forever. Grim trigger is a severe trigger strategy since a single defection brings about an eternal end to cooperation, in contrast to the much more forgiving tit for tat.

**Pareto:** Pareto was an Italian economist who lived from 1848 to 1923. He argued that an individual's preferences were the beginning point of economic analysis, and only ordinal and not cardinal payoffs were important. Keeping with this, he developed the notion of a Pareto optimal outcome in which no member of society can be made better off without hurting, or decreasing the payoffs of someone else.

**Pareto efficiency:** Named after Vilfredo Pareto, Pareto efficiency (or Pareto optimality) is a measure of efficiency. An outcome of a game is Pareto efficient if there is no other outcome that makes every player at least as well off and at least one player strictly better off. That is, a Pareto Optimal outcome cannot be improved upon without hurting at least one player. Often, a Nash Equilibrium is not Pareto efficient implying that the players' payoffs can all be increased.

**Pareto optimal:** Pareto optimality is a measure of efficiency. An outcome of a game is Pareto optimal if there is no other outcome that makes every player at least as well off and at least one player strictly better off. That is, a Pareto Optimal outcome cannot be improved upon without hurting at least one player. Often, a Nash Equilibrium is not Pareto Optimal implying that the players' payoffs can all be increased.

**Pareto dominated:** An outcome of a game is Pareto dominated if some other outcome would make at least one player better off without hurting any other player. That is, some other outcome is weakly preferred by all players and strictly preferred by at least one player.

**Pay-off**: Payoffs are numbers which represent the motivations of players. Payoffs may represent profit, quantity, "utility," or other continuous measures (cardinal payoffs), or may simply rank the desirability of outcomes (ordinal payoffs). In all cases, the payoffs must reflect the motivations of the particular player.

**Player:** Any participant in a game who has a nontrivial set of strategies (more than one) and selects among the strategies based on payoffs.

**Parallel Subgame Perfect Equilibrium (PSPE):** It is an equilibrium in which the player’s strategy constitutes Nash equilibrium in every parallel subgame set of the original game which is equal to the appropriate response, because it is determined by the maximization of the minimum pay-offs under consideration of all SPE strategies in a time period.

**Pathway:** Every set of edges in extensive form of the game or game tree, which begins from the first node in the tree(root) and eventually arrives at the repetition stages.

**Rationality:** One of the most common assumptions made in game theory (along with common knowledge of rationality). In its mildest form, rationality implies that every player is motivated by maximizing his own payoff. In a stricter sense, it implies that every player always maximizes his utility, thus being able to perfectly calculate the probabilistic result of every action.

**Strategic form:** The strategic (or normal) form is a matrix representation of a simultaneous game. For two players, one is the "row" player, and the other, the "column" player. Each rows or column represents a strategy and each box represents the payoffs to each player for every combination of strategies. Generally, such games are solved using the concept of a Nash equilibrium.

**Strategy:** A strategy defines a set of moves or actions a player will follow in a given game. A strategy must be complete, defining an action in every contingency, including those that may not be attainable in equilibrium.

**Subgame:** A subset or piece of a sequential game beginning at some node such that each player knows every action of the players that moved before him at every point. Subgame perfect equilibria discovered by backward induction are Nash equilibria of every subgame.

**Subgame perfect equilibrium(SPE):** A subgame perfect Nash equilibrium is an equilibrium such that players' strategies constitute a Nash equilibrium in every subgame of the original game. It may be found by backward induction, an iterative process for solving finite extensive form or sequential games. First, one determines the optimal strategy of the player who makes the last move of the game. Then, the optimal action of the next-to-last moving player is determined taking the last player's action as given. The process continues in this way backwards in time until all players' actions have been determined.

**Tit for Tat:** A type of trigger strategy usually applied to the repeated Prisoner's Dilemma in which a player responds in one period with the same action her opponent used in the last period.

**Zero sum game:** A zero sum game is a special case of a constant sum game in which all outcomes involve a sum of all player's payoffs of 0. Hence, a gain for one participant is always at the expense of another, such as in most sporting events. Given the conflicting interests, the equilibrium of such games is often in mixed strategies.

**A-H:** *Mtb*'s strategies ($S_A$-$S_H$) or final outcomes begin the game (first node of the game) in the game tree (figure1). Thus, pathways called A-H in the text based on this origination.
**$S_A$-$S_H$:** *Mtb* Strategy
**$S_a$-$S_h$:** Host Strategy
**$n_0$:** The number of unrepeated parallel subgames.
**$n_r$:** The number of repeated parallel subgames that passed after the number of repetition times "t".
**x:** The number of strategies which are matched with the PSPE in unrepeated parallel subgames.
**y:** The number of strategies which are unmatched with the PSPE in unrepeated parallel subgames.
**x':** The number of strategies which are matched with the PSPE in repeated parallel subgames.
**y':** The number of strategies which are unmatched with the PSPE in repeated parallel subgames.
